## Supplemental Information for "Remodeling of lipid-foam prototissues by network-wide tension fluctuations induced by active particles"

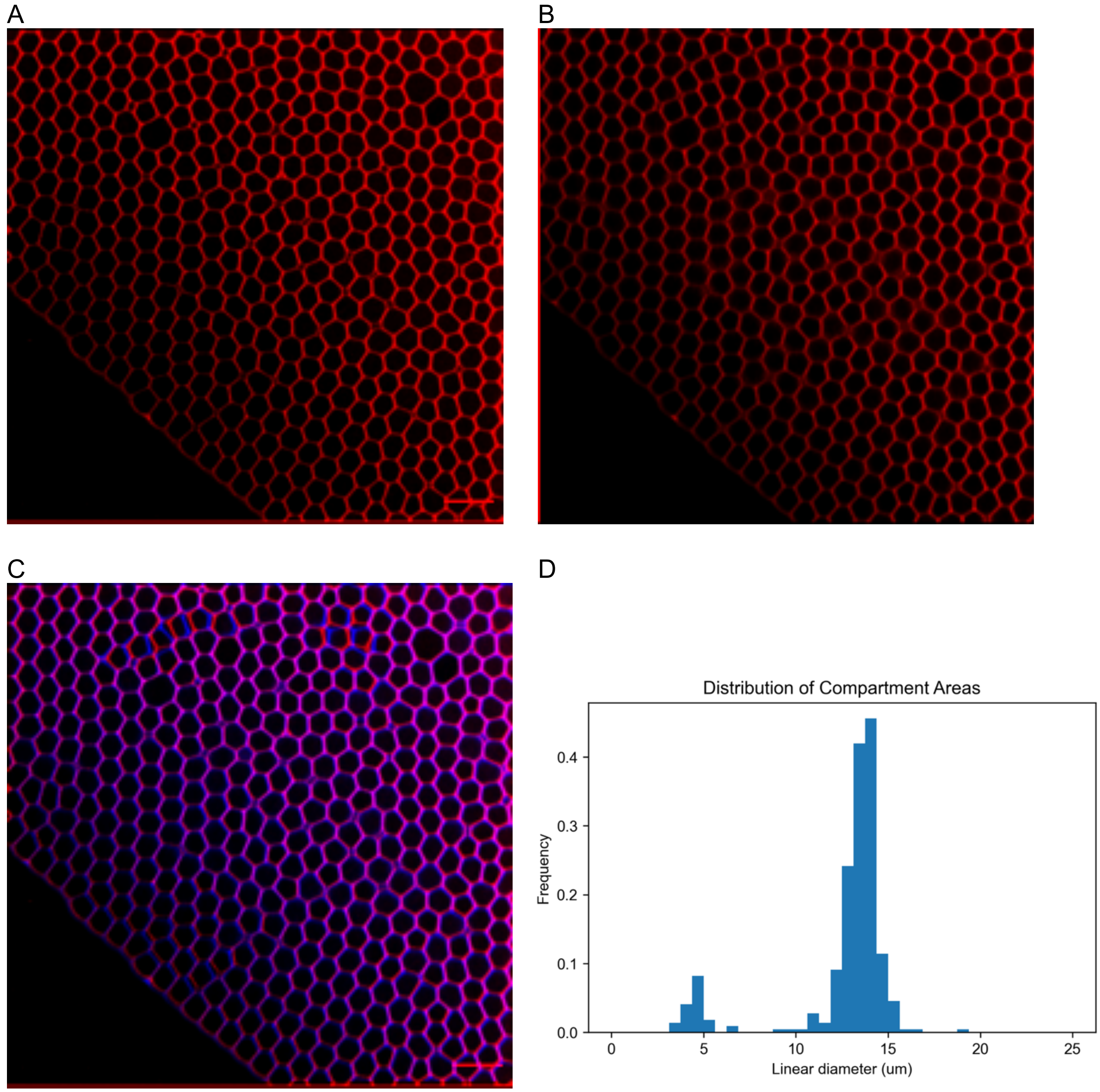


Supplementary Figure 1 – Stability of the structure at time A) t = 0 B) t = 44 h and C) overlay in red (t =0) and blue (t = 44 h). D) Histrogram corresponding to panel A. Scale bar 100 μm.

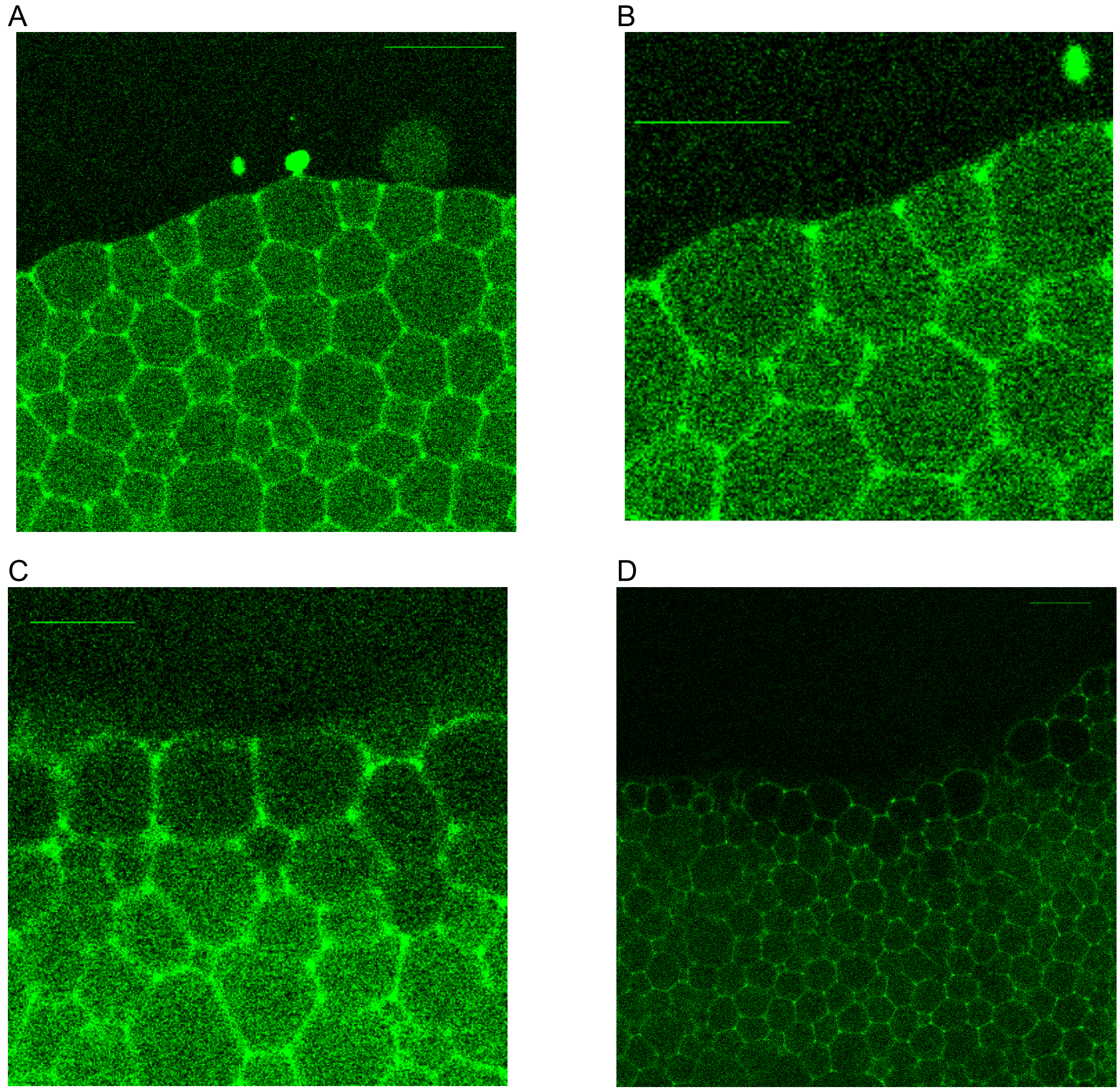


Supplementary Figure 2 – A) Green signal shows water-soluble fluorescein encapsulated. Signal along lipid bilayers is bleed-through from membrane dye. Scale bar: 50 µm. B) Same image with zoom on interface region shows that compartments homogeneously encapsulate water-soluble dye. Scale bar: 30 µm. C) Alpha-hemolysin (αHL) was added at a concentration of 6.7 ng/µL and the structure was imaged 5 minutes after addition. Permeation of the dye in the compartments facing the outside water phase can be seen. Scale bar: 30 µm. D) 30 minutes after addition of αHL. Scale bar: 50 µm.

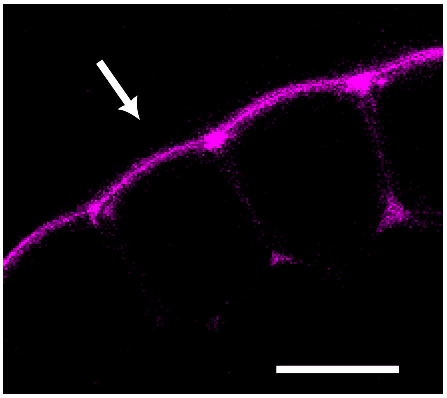

Supplementary Figure 3 – Penetration of water soluble but hydrophobic dye (Atto 647N) added from the direction indicated by the arrow. Over time, the dye concentrates in the lipid bilayer / vertex structure and diffuses along the hydrophobic continuous phase. Scale bar 20 μm.

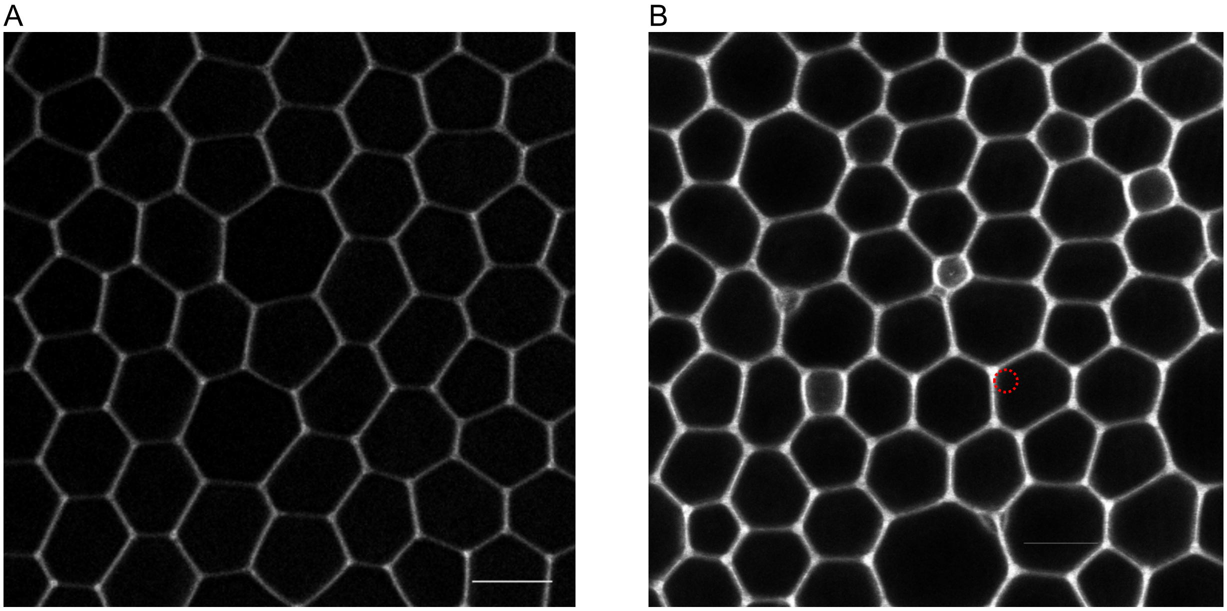


Supplementary Figure 4 – Decorated vertices with optically resolvable oil pockets. A) Typical structure shows no optically resolvable curvature at the vertices. B) Some preparations show excess oil or lipids (variations in grayscale contrast) and optically resolvable curvature (indicated by red circle) at the vertices/edges.

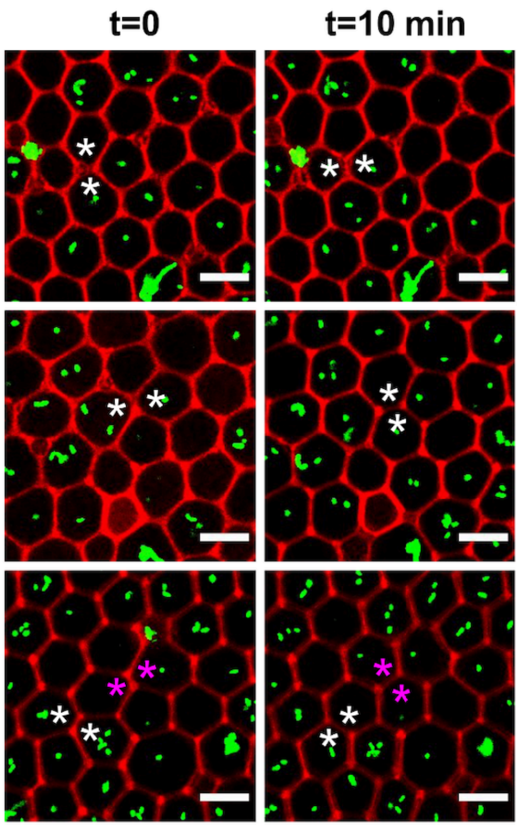


Supplementary Figure 5 – Additional examples of T1 transitions (stars) observed when bacteria were encapsulated. Scale bar 20 μm.

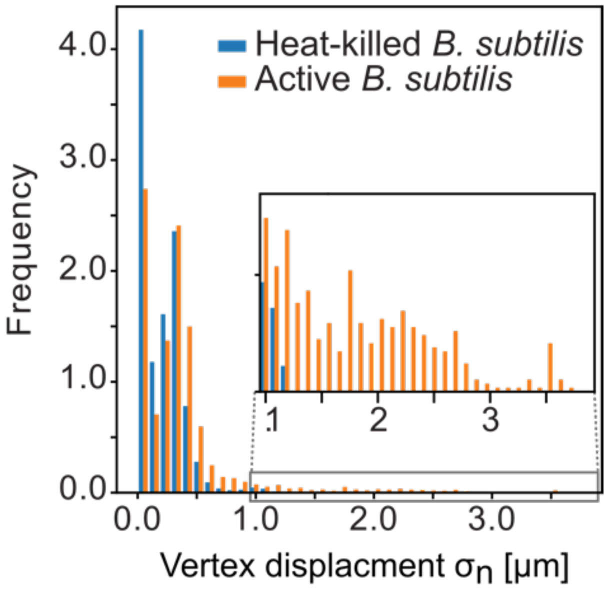


Supplementary Figure 6 - Histogram of vertex displacement obtained for active and heat-killed bacteria. Inset shows enlarged view of the indicated region. We found that vertices of compartments filled with live bacteria showed larger displacements then heat-killed, non-swimming, bacteria. Averaging 600 frames and 68 vertex locations over 10 minutes, we found an average standard deviation in vertex position of 0.2 µm when heat-killed, passive, bacteria were encapsuled. The average standard deviation in vertex position was found to be 0.37 µm when live, swimming bacteria were encapsulated, a significant increase compared to that of heat-killed bacteria. Notably, the distribution of average displacements was not Gaussian; instead, a long tail of displacement values up to 3.5 µm were found with swimming bacteria (inset).

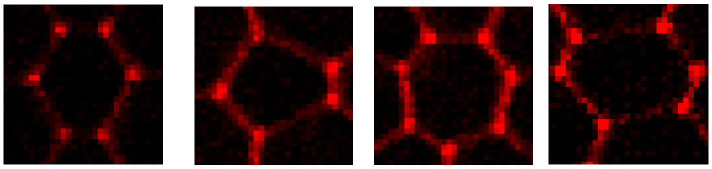

Supplementary Figure 7 – Examples images of non-hexagonal compartment shapes. Stochastic variations of the abundance of such compartments in each preparation lead to variations in shape index between samples.

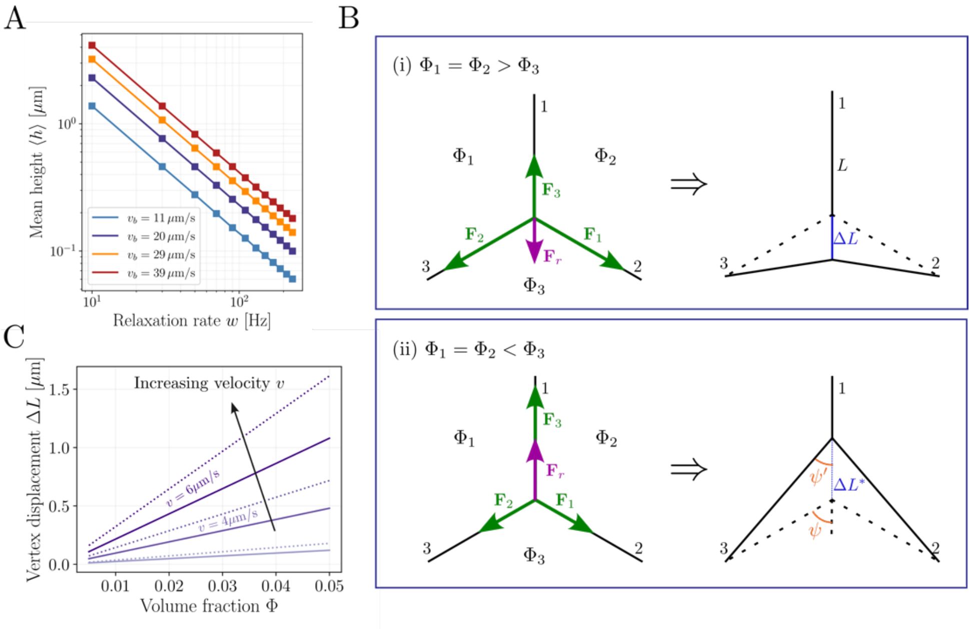


Supplementary Figure 8 – Minimal model on membrane height deformations and vertex displacements. (A) Steady-state prediction of the membrane height (square markers) matches well with the numerically obtained values for the mean height <*h*> (solid lines). The mean height decreases with the inverse of the relaxation rate *w* for different choices of swimming velocities (different colors). (B) Illustration of possible displacements of a single vertex driven by the activity of bacteria in the surrounding compartments. (Top) The case of compartments 1 and 2 having an equal density of bacteria that is larger than that of compartment 3. Each compartment generates a force $\mathbf{F}_i$ which can be projected onto the vertex, acting along the edge complementary to the compartment. As $|\mathbf{F}_1 + \mathbf{F}_2| > |\mathbf{F}_3|$, the resultant force $|\mathbf{F}_r|$ points along edge 1 by extending it, leading to a vertex displacement of ΔL. (Bottom) The case of compartments 1 and 2 having an equal density of bacteria that is smaller than that of compartment 3. As $|\mathbf{F}_1 + \mathbf{F}_2| < |\mathbf{F}_3|$, the resultant force $|\mathbf{F}_r|$ points along edge 1 by contracting it, leading to a vertex displacement of $\Delta L^*$. The angles $\psi$ and $\psi'$ denote half of the vertex angle before and after displacement, respectively. C) Scaling of the vertex displacements ΔL as a function of directional volume fraction $\Phi$ for different values of bacteria velocities *v* (color coded). Dotted lines represent vertex displacements $\Delta L^*$ that correspond to an edge contraction event from an angle $\psi = 60°$ to $\psi' = 45°$, which leads to larger deviations from ΔL as the velocity increases.

**Supplementary Note 1**

We calculated the Span-80 concentration in the oil phase as 5.9 mM which is known from the transferred volume during the preparation. The oil phase was in contact with a water phase of equal volume over the typical experimental timescales of many hours, suggesting that the Span-80 will have equilibrated between the phases. With the partition coefficient of Span-80 between octanol and water being logP = 6 (PubChem CID 9920342), we estimate a Span-80 concentration to be 6 nM in the water phase. If equilibration would not be reached the concentrate of Span-80 in the water phase would be even lower. We estimated the amount of lipids in the water phase from the total size of the structure. Taking the typical diameter of a single compartment is R = 20 µm, we can calculate the number of lipid area per compartment as $4\pi R/A$ with $A$ representing a area per lipid of 0.5 nm². Together with a conservative estimate of a million compartments (as judged by the typical area occupied on a 96-well plate), this gives a lipid concentration in excess of 100 nM. Comparing to the maximal concentration of 6 nM of Span-80 this shows that Span-80 can be considered a minor impurity compared to the phospholipids.

**Supplementary Note 2**

To further expand our quantitative understanding of the chemical-mechanical coupling between swimming bacteria and deformations in the tissue mimetic, we consider a simplified one-dimensional lattice mode with elastic coupling between *N* vertices with displacement $u_j$. As the measured $l_{corr}$ indicates that only nearest neighbors contribute to the coupling, we calculated the power spectrum of the Fourier transform $\mathcal{F}(u_0)$ for *N* = 2, which we then compare to the experimentally obtained power spectra.

$$(1)\ |\mathcal{F}(u_0)|^2 = \frac{2k_bT_{eff}}{b(f^2+2f_0^2)}$$

With the cross-over frequency $f_0 = \frac{k}{b}$, stiffness of the elastic coupling $k$, drag coefficient $b$, and (effective) temperature $T_{eff}$. Introduction of an effective temperature allows us to capture the effect of the bacteria collisions, which, on sufficiently long timescales, appear as a random (uniformly distributed) force, a standard approach for systems with active particles. In the experiment, both alive and heat-killed bacteria groups showed a comparable spectrum to Eq. (2). Consistent with the previous analysis live bacteria show higher PSD values, corresponding to larger vertex displacement, than the head-killed control. For both samples some deviations towards higher frequencies were observed (Fit parameters are in caption of Fig. 3). The high-frequency deviations might stem from noise of the vertex detection which leads to leveling off for small sub-nanometer displacements. Additionally, at higher frequencies, deviations are expected as the effective temperature approximation is not valid anymore. Nevertheless, the approximate agreement between Eq. (2) and experiments allowed us to establish that (1) The vertices indeed form an elastic lattice with coupling provided by the lipid bilayers, corroborating the model for cooperative tension generation suggested above, (2) The dynamics of the tissue mimetic were relatively slow with $f_0 \sim 10^{-2}$ Hz , with a large drag coefficient $b \sim 0.5$ µNs/m. These values were comparable to relaxation timescales found in in-vivo tissue measurements in the order of $10^{-1} - 10^{-2}$ Hz (Ref.[36]) and (3) The increase in effective temperature of $\frac{T_{eff}}{T} = 6.5$ by the encapsulated bacteria is comparable in magnitude found in biological systems. For example, ATP-driven membrane fluctuations in red blood cells result in $\frac{T_{eff}}{T} = 3$,[37,38] underlining the fact that the high densities of bacteria encapsulated here provide a strong non-equilibrium driving force. Taken together, these results demonstrate the tissue mimetic can be described as an elastic lattice that shows similar dynamics as biological tissues.

### Supplementary Methods

*Theoretical description of membrane deformations driven by bacteria.*

To understand the dynamics of vertex fluctuations, we first wanted to clarify whether the influence of bacteria collisions with a given membrane patch could lead to off-plane displacements of the membrane. For this, we first turned to a minimal theoretical description using Monge representation of the membrane from a relaxed reference plane, and denoted the orthogonal displacements from this relaxed state by the height function *h(x,t)*. In addition, to have an effective description of bacterial activity during its contact with the membrane, we'll omit "on / off" rates of bacteria and simply focus on the active displacements induced by their swimming against the membrane with an overall effective "occupation density" of bacteria b(x,t). Because bacteria induce local deformations on the membrane height via their active swim velocity $v_b$, this interaction bears similarities with membrane-protein complexes where active particles on the membrane act to locally shape its geometry through nonequilibrium forces. Following the theoretical framework in [Cagnetta et al PRL 2018], we can thus write for the time evolution of the membrane height function:

$$\frac{\partial h}{\partial t} = D\,\frac{\partial^2 h}{\partial x^2} - wh + \lambda b\left(1 - \left(\frac{\partial h}{\partial x}\right)^2\right), \quad \text{(S1)}$$

where the first term describes thermal fluctuations of the height with an effective diffusion coefficient *D*, the second term accounts for the membrane relaxation due to bending modulus $\kappa$ and tension $\sigma$ , defined by the Fourier transformed relaxation rate $w(k) = \Lambda_k(\kappa k^4 + \sigma k^2)$. Here the Oseen tensor in Fourier space is given by $\Lambda_k = (4\eta k)^{-1}$ [Turlier and Betz 2018]. The effect of bacterial activity with speed $v_b$ on the membrane is described by the last term, where the parameter $\lambda$ connects the bacteria density to the "forward velocity" of local deformations induced by swimming via $v_b = \lambda b$ [Gov PRL 2006].

We can now complement this with the evolution for the bacterial occupation density on the membrane with a simple diffusion-advection equation: $\frac{\partial b}{\partial t} = D_b\,\frac{\partial^2 b}{\partial x^2} - \chi\frac{\partial}{\partial x}\left(b\,\frac{\partial h}{\partial x}\right)$, (S2)
where $D_b$ represent the effective diffusion of the bacteria along the membrane, while the second term defines an advective flux through curved regions of the membrane patch with strength controlled by $\chi$ . This term thus provides a minimal curvature-activity coupling that also effectively accounts for the fact that while pushing the membrane, the bacteria dwell for a certain amount of time locally in contact with the membrane, before "tumbling away" from the membrane patch into the bulk of the vesicle.

Interestingly, Eq. (S1) already shows a simple estimate for the mean displacements of the membrane in the limit of small local curvatures $\partial h/\partial x \sim 0$ . It predicts that the steady-state height will be determined by the product of bacterial velocity $v_b$ and membrane relaxation time $\tau^{-1} = w$:

$$h^{st} = v_b\tau, \qquad \text{(S3)}$$

A membrane patch of the order $L\ \cong 15\mu m$ , which corresponds to the average edge length between two vertices in the experiment, indicates a mode of approximately $k\ = 2\pi/L \cong$

$0.42\ \mu m^{-1}$. Using a bending modulus of around $\kappa = 20\ k_B T$ with $T = 300K$ , a surface tension of around $\sigma \cong 10^{-3} N/m$ as determined from the experiments, and a viscosity of $\eta \cong 10^{-3} Pa\ s$, we get for the membrane relaxation rate $w(k) = (\kappa k^3 + \sigma k)/4\eta \cong 10^5 s^{-1}$. For experimentally obtained values of $v_b \sim 10 \mu m/s$ (see main text), this relaxation rate then indicates a mean displacement of the order $h^{st} \sim 0.1 nm$, suggesting that the membrane patch between two vertices of the network is essentially flat. For membrane tension values of the order $\sigma \sim 10^{-6} - 10^{-7} N/m$, on the other hand, the same parameter set would lead to a mean membrane displacement of around $h^{st} \sim 0.4 \mu m$, which highlights the key role of membrane tension in its mechanical coupling with an active bath. To test if the steady state approximation given in Eq.(S3) holds robustly for different parameter choices, we numerically solved the coupled PDE system using different values of bacterial activity fixed by $b_0 \equiv b(t = 0)$ (using the Python package py-pde [Zwicker, Journal of Open Source Software 2020]). We found that $h^{st}$ robustly predicted the long-time average height values obtained while changing the relaxation rate *w* and the active velocity $v_b$, see Fig.S5A.

*Minimal model for vertex displacements driven by bacterial activity.*

Having found that each edge of the lipid network exhibits only nano-scale deformations due to the rather large membrane tension, we then turned to a toy model to describe the force balance conditions at each vertex of the network. We first assumed that in each compartment bacteria on average exert an equal swimming pressure on each edge, and we argued that this would lead to a net force projected onto the vertices of the compartment. Therefore, we can for simplicity focus on the force balance condition at a single (three-valent) vertex.

Let us now denote by $\Phi_i$ the volume fraction of bacteria in each of the three compartments surrounding a vertex labelled with *i*=1,2,3. We can likewise label each edge adjacent to the vertex with indices *j*=1,2,3. As illustrated in Fig.S5B, the swimming force $\mathbf{F}_1$ generated in compartment 1 will then be directed along the edge 2, the force $\mathbf{F}_2$ along the edge 3, and the force $\mathbf{F}_3$ along the edge 1. The resultant force on the vertex will then be given by the sum $\mathbf{F}_r = \Sigma_j \mathbf{F}_j$. For a single compartment, the collective pressure of bacteria arising from their active swimming takes the form [Xie et al PRL 2022]:

$P_i = \frac{3\eta\tau}{4R^2} v^2 \Phi_i$ ,        (S4)

where $\eta$ is the viscosity, *R* is the effective radius, $\tau$ is the persistence time, *v* is the swim velocity of bacteria. The vertex displacement from its thermal equilibrium position has to be balanced by elastic forces generated through edge length variations due to this swimming pressure. Each compartment will then induce a net swimming force of magnitude $|\mathbf{F}_i| = P_i A$, where *A* is an effective membrane area corresponding to a single edge (i.e. including the height in z-direction). In general, the resultant force vector can have arbitrary directions, as the distribution of bacteria densities $\Phi_i$ in all three compartments surrounding the vertex need not be identical. However, if we assume a homogeneous distribution of bacteria within compartments, the resultant force should point at the opposite direction as the vectorial volume fraction $\mathbf{\Phi} = \mathbf{\Phi}_1 + \mathbf{\Phi}_2 + \mathbf{\Phi}_3$ with a magnitude $F_r$ proportional to $\Phi = |\mathbf{\Phi}|$. Here, $\mathbf{\Phi}_i$ are vectors of length $\Phi_i$ pointing at the center-of-mass of the *i*-th compartment. We now argue that the magnitude of the resultant force should be related to the swimming pressure of each compartment by

$F_r = \frac{3\eta\tau A}{4R^2} v^2 \Phi$.        (S5)

Let us now for simplicity focus on two minimal examples to explain the elastic force balance condition and to link vertex displacements with the mechanical properties of the membrane:

(i) **Two compartments winning against one compartment:** For $\Phi_1 = \Phi_2 > \Phi_3$, where the compartments 1 and 2 have an equal density of bacteria which is larger than that of the compartment 3 (see top panel in Fig.S5B), the resultant force will act along the edge 1 to extend it by an amount Δ*L*. Assuming that only edge extension induce elastic forces, i.e. that the compression modes are negligible as the membrane is fluid, the new vertex position will then be determined by a force balance condition:

$F_r = \sigma_1 \Delta L$ ,     (S6)

where $\sigma_1$is the tension corresponding to edge 1. Using Eq.(S5), this suggests that the vertex will be displaced from its "thermal" (i.e. passive) position by an amount:

$\Delta L = \frac{3\eta\tau A}{4R^2}\frac{v^2\Phi}{\sigma_1}$ .     (S7)

(ii) **One compartment winning against two compartments:** For $\Phi_1 = \Phi_2 < \Phi_3$, the resultant force will again act along the edge 3 but now to compress it by an amount Δ*L*, while extending the edges 2 and 3 (see bottom panel in Fig.S5B). Because in this configuration the stretched edges 2 and 3 will partially act as two springs connected in parallel, we can use a simple trigonometric relation to determine the vertex position from the force balance condition as:

$F_r \cong (\sigma_2 + \sigma_3)\Delta L \cos(\psi)\cos(\psi')$ ,     (S8)

where $\psi$ and $\psi'$ are half of the vertex angle between edges 2 and 3 before and after displacement, respectively. For angles close to $\psi = 60°$ and $\psi' = 45°$ (e.g. a vertex angle of $120°$ from a hexagonal lattice will be reduced to an angle $90°$ ), and for identical edge tensions $\sigma_2 = \sigma_3 = \sigma$, this indicates that the resultant force will in general be $F_r = \gamma\sigma\Delta L$ with $\gamma \cong 0.7$ . The corresponding vertex displacement governed by the force balance condition will then be:

$\Delta L^* = \frac{3\eta\tau A}{4R^2}\frac{v^2\Phi}{\gamma\sigma}$ .     (S9)

For deformations from the reference state of a hexagonal compartment, the prefactor $\gamma$ will in general take values between 0.5 and 1. The relations given in Eqs.(S7&S9) allow us to determine the linear scaling of the vertex displacement as a function of the directional volume fraction Φ after inferring the remaining parameters from experimental data, see Fig.S5C for different vertex displacement values e.g. for different bacterial velocity *v* values. Note that, depending on the configuration, the observed vertex displacement will have a spread simply arising from mechanical constraints.

*Calculation of the power spectrum*

The tension acting on vertex *j* is given by a force balance as $T_j = k(2u_j - u_{j+1} - u_{j-1})$, where $u_j$ is the vertex displacement from its resting position and $k$ is the stiffness of the elastic coupling between vertices. To study the dynamics of the system, we considered an overdamped Langevin equation where every vertex is subject to stochastic agitation by a diffusion term $\eta(t)$ with friction coefficient $b$. Consistent with the low Reynolds number in our experimental system, we balanced motion of the vertices with a damping coefficient $b$ and the tension $T_j$, resulting in the equation $b\frac{du_j}{dt} = k(2u_j - u_{j+1} - u_{j-1})$, with a time constant $\tau = \frac{b}{k}$. To study the dynamics of the system, we considered an overdamped Langevin equation where every vertex is subject to stochastic agitation by a white noise diffusion term $\eta(t)$.

(1) $b\frac{du_j}{dt} = k\left(2u_j - u_{j+1} - u_{j-1}\right) + \eta(t)$

The power spectrum was calculated by taking the Fourier transform

(2) $-bi\omega t\, U_j(\omega) = k\left(2U_j(\omega) - U_{j+1}(\omega) - U_{j-1}(\omega)\right) + \Xi(\omega)$

For a lattice N=2 two linear equations with two unknows $U_0$ and $U_1$ are obtained. The resulting power spectrum $|U_0(\omega)|^2$ shown in eq. (2) was then calculated with $\Xi(\omega) * \bar{\Xi}(\omega) = k_B T$.
